# BBB-on-a-chip with Integrated micro-TEER for permeability evaluation of multi-functionalized gold nanorods against Alzheimer’s disease

**DOI:** 10.1101/2022.09.05.505851

**Authors:** Sujey Palma-Florez, Adrián López-Canosa, Francisco Morales-Zavala, Oscar Castaño, M.J. Kogan, Josep Samitier, Anna Lagunas, Mónica Mir

## Abstract

**Background:** The lack of predictive models that mimic the blood-brain barrier (BBB) hinders the development of effective drugs for neurodegenerative diseases. Animal models behave differently from humans, are expensive and have ethical constraints. Organ-on-a-chip (OoC) platforms offer several advantages to resembling physiological and pathological conditions in a versatile, reproducible, and animal-free manner. In addition, OoC give us the possibility to incorporate sensors to determine cell culture features such as trans-endothelial electrical resistance (TEER). Here, we developed a BBB-on-a-chip (BBB-oC) platform with a TEER measurement system in close distance to the barrier used for the first time for the evaluation of the permeability performance of GNR-PEG-Ang2/D1 for Alzheimer’s disease. GNR-PEG-Ang2/D1 is a therapeutic nanosystem previously developed by us consisting of gold nanorods (GNR) functionalized with polyethylene glycol (PEG), angiopep-2 peptide (Ang2) to overcome the BBB and the D1 peptide as beta amyloid fibrillation inhibitor, finally obtaining GNR-PEG-Ang2/D1 which showed to be useful for disaggregation of the amyloid in *in vitro* and *in vivo* models. In this work, we evaluated its cytotoxicity, permeability, and some indications of its impact on the brain endothelium by employing an animal-free device based on neurovascular human cells.

**Results:** In this work, we fabricated a BBB-oC with human astrocytes, pericytes and endothelial cells and a TEER measuring system (TEER-BBB-oC) integrated at a micrometric distance of the endothelial barrier. The characterization displayed a neurovascular network and the expression of tight junctions in the endothelium. We produced GNR-PEG-Ang2/D1 and determined its non-cytotoxic range (0,05–0,4 nM) for plated cells included in the BBB-oC and confirmed its harmless effect at the highest concentration (0.4 nM) in the microfluidic device. The permeability assays revealed that GNR-PEG-Ang2/D1 cross the BBB and this entry is facilitated by Ang2 peptide. Parallel to the permeability analysis of GNR-PEG-Ang2/D1, an interesting behavior of the TJs expression was observed after its administration probably related to the ligands on the nanoparticle surface.

**Conclusion:** BBB-oC with TEER integrated setup was proven as a functional and throughput platform to evaluate the brain permeability performance of nanotherapeutics in a physiological environment with human cells, putting forward a viable alternative to animal experimentation.

## BACKGROUND

Neurodegenerative diseases (NDDs) represent a major threat to the health of the population. In recent years, these age-dependent diseases have become more prevalent, partly because life expectancy has increased [1]. Examples of NDDs are Alzheimer’s disease (AD), Parkinson’s disease (PD), Huntington’s disease (HD), and amyotrophic lateral sclerosis (ALS). Unfortunately, the drugs intended to reach the central nervous system (CNS) have much higher failure rates preclinically and clinically than non-CNS drugs, and the worst outcomes are observed in chronic NDDs such as AD, PD, ALS, and neuromuscular disorders [2]. Particularly in AD, there are no effective treatments available and more than 200 therapeutic agents in evaluation programs have failed or deserted [3–5].

One of the reasons for the clinical failure of CNS-drugs is the brain anatomy itself. The brain is a protected organ by the blood-brain barrier (BBB), one of the most restrictive barriers in the body which acts as a natural guard [6–8] avoiding the entrance of neurotoxic agents, invasive pathogens, and most of therapeutic drugs [9]. The impermeability of the BBB is mainly because of the tight junctions (TJs) assembled between brain endothelial cells (EC) which are formed by structural proteins such as occludins, claudins, and junctional adhesion molecules [10] that regulate the passage across the cerebral endothelium.

Another reason is the lack of prediction offered by the *in vivo* models due to the physiological differences between humans and animals. In recent years, animal models have been genetically manipulated to recreate some human diseases, however, chronic NDDs models have rarely proven to be predictive [2]. In addition, these models are expensive, ethically questionable, and frequently inadequate to understanding at the molecular/cellular level the complexity of the disease [11,12].

Organ-on-a-chip (OoC) is an attractive emerging technology that overcomes several drawbacks regarding animal models. OoC has a versatile design, low cost and it is an animal-free approach to mimic *in vivo* physiological and pathological conditions [13] for the study of drug permeability and efficacy, and disease progression [14]. Recently, several OoC devices have been developed to mimic biological barriers as the BBB (BBB-oC). Nonetheless, several BBB-oC platforms have been shown no physiological relevance as they exhibit permeability coefficient for standard molecules such as dextran (70 kDa), in the order of 10^−5^, which is clearly above *in vivo* values around 10^−8^ [15]. Some reasons to explain this difference are the use of non-human cells to assemble the neurovascular network, as well as the absence of other key cell types such as astrocytes, pericytes and microglia which supplying promoting factors for the proper development of the brain endothelium [16,17]. On the other hand, factors related to the microdevice setup also influence the correct maturation of the endothelium such as the use of commercial membranes for the cell seeding like, polycarbonate membrane, which are too wide and therefore hinder the cell-to-cell contact [18–20]. One strategy to overcome this problem, is the use of 3D extracellular matrix (ECM) such as collagen or fibrinogen which act as a natural scaffold for cell assembly allowing direct cell-to-cell and cell-to-matrix interaction [21,22].

Furthermore, detection platforms can be incorporated into BBB-oC to monitor features as barrier permeability, drug efficacy or toxicity. Trans-endothelial electrical resistance (TEER) is a widely used quantitative method for assessing the barrier integrity and functionality by the measurement of the electrical resistance across the cellular monolayer. BBB-oC reported have shown TEER values around 3000 and 20000 Ω.cm^2^ which are close to healthy human BBB (1500 and 8000 Ω.cm^2^) [18,19,23–25], however, usually the measurement setup is still not optimized and leads to miss estimation of TEER results. Some important aspects to consider for the correct TEER read-out are the electrodes position respecting the endothelial cells. Several previous works placed electrodes far away from the endothelial barrier causing higher resistance contribution from other elements such as the cell medium solution or the commercial membranes. Additionally, factors as the ratio between the electrode and endothelial barrier size is crucial to generate a homogeneous current density through the BBB to achieve a proper measurement [26–28].

Since AD is characterized by neuronal death associated partly with extracellular accumulation of beta amyloid (Aβ) plaques, several therapeutics agents are focused on altering the Aβ fibrillation process and promote their clearance [29,30], although none have been fully successful. Nanotechnology is a rising field with high potential to offer new opportunities for AD treatment and diagnosis. A previous work assembled a biologically inspired nanostructure (ApoE3-rHDL) able to pass through the BBB, target Aβ oligomers and promote the degradation of the Aβ structure through the lysosomal pathway *in vivo* [31]. As well, gold nanoparticles previously showed to trap Aβ monomers to suppress the fibril formation [32]. Specifically, our group developed a nanosystem against AD consisting of gold nanorods (GNR) coated with polyethylene glycol (PEG) and biofunctionalized with D1 peptide that acts as Aβ sheet breaker (D1) and Angiopep-2 (Ang2) peptide to enhance its transport through the BBB. D1 peptide with sequence qshyrhispaqv contains d-amino acids [33] and has a high affinity for Aβ monomers and oligomers avoiding Aβ aggregation [34–36]. This peptide has been discovered by a phage display scanning and the presence of d-amino acids increase the stability in the biological medium. On the other hand, Ang2 is a peptide that binds to low-density lipoprotein receptor-related protein 1 (LRP1) present in the brain endothelium allowing it to cross the BBB. Our results revealed that the nanosystem (GNR-PEG-Ang2/D1) was able to inhibit Aβ growth *in vitro* and decrease the toxicity of Aβ aggregates in a *Caenorhabditis elegans in vivo* model [37]. Recently, this nanosystem demonstrated be helpful for AD diagnosis through the detection of Aβ by computerized tomography and Aβ disaggregation in mice [38]. However, despite the great potential of nanotechnology, few works have used BBB-oC for the evaluation of nanoparticles permeability [39–41], but to our knowledge, none has yet tested any therapeutic nanoparticle as a potential treatment against NDDs.

In this work, we present an animal-free BBB-oC model consisting of human endothelial cells in co-culture with human astrocytes and pericytes cells assembled in a 3D ECM scaffold. A micro-TEER measuring system was integrated considering critical aspects such as the position and dimensions of the electrode, placing the TEER in a micrometric distance from the barrier for the correct read-out of TEER to monitoring the endothelium maturation. Finally, we evaluated the cytotoxicity and permeability performance of GNR-PEG-Ang2/D1.

## MATERIALS AND METHODS

### Materials

All reagents were purchased from Sigma-Aldrich (MO, USA) unless otherwise indicated.

Human hippocampal astrocytes, human brain-vascular pericytes, poly-L-lysine, trypsin/EDTA 0.05% and astrocyte and pericyte media were supplied by Sciencell (CA, USA). Human brain endothelial cell line (hCMEC/D3) and EndoGRO™ MV cell medium kit were purchased from Merck-Millipore (MA, USA). Trypsin/EDTA 0,25%, calcein-AM (2µM), ethidium homodimer-1 (4µM), DAPI, Mo-Hu monoclonal ZO1 antibody (0.5 mg/mL), Hoetch 33342 (10 mg/mL), secondary antibodies anti-rabbit Alexa 568 (2 mg/mL) and anti-mouse Alexa 488 (2 mg/mL), and Alexa 647 hydrazide (1 mg) were obtained from Thermo Fischer Scientific (MA, USA). Rab-Hu polyclonal VE-cadherin (1mg/mL) and Rab-Hu monoclonal LRP1 (0.46 mg/mL) antibodies and phalloidin-iFluor 594 reagent were supplied from Abcam (Cambridge, UK). Annexin-V was provided by BioVision (CA, USA). Staurosporine was purchased from Selleckhem (Madrid, Spain).

HS-PEG-OMe and HS-PEG-COOH MW 5000 Da were provided from JenKem Technology (TX, USA). Angiopep-2 and D1 peptides were synthesized through solid-phase synthesis technique at the laboratory of Nanotechnology at Universidad Andrés Bello (Santiago, Chile).

The resins SU8-2100 and AZ 5214, and AZ 100 remover were obtained from Microchemicals (Ulm, Germany). The acetate masks were fabricated by JD Photo Data (UK) and the glass coverslip (76 mm x 26 mm x 1.5 mm) were supplied from Menzel-gläser (Braunschweig, Germany). The SYLGARD™ 184 Silicone Elastomer Kit for polydimethyl siloxane (PDMS) was purchased from Ellsworth (WI, USA). Deionized Milli-Q water of 18 MW·cm was obtained through Milli-Q® purification equipment from Meck Millipore (Darmstadt, Germany) using previously deionized conventional water (acid and basic columns).

### Fabrication of the microfluidic device

The microfluidic device was designed with AUTOCAD software (AUTODESK, USA) and the design was printed as an acetate mask. Master molds were obtained in 100 mm diameter silicon wafers using standard photolithography techniques in a clean room environment. Then, the 100 mm silicon wafers were activated in the plasma chamber at high mode (10.5 W) for 1 min and two layers of SU8-2100 photoresist were spun to a total height of 120μm. Then, the photoresist was soft baked for 25 min and exposed using an I-line mask aligner to transfer the design from the acetate mask. After that, the wafers were post-baked, developed for 20 min and then hard baked for 30 min at 95°C and 10 min at 65°C. Finally, the wafers were silanized for 1 h with trifluorosilane. The PDMS replicas were obtained using the well-established method of soft lithography [42]. PDMS was mixed in a ratio 10:1 (base:crosslinking agent), poured into the mold and cured for 2 h at 65°C. Every single device was cut and bonded with a glass slide through irreversible plasma treatment for 30 s at high mode (10.5 W). Finally, the microdevices were left in the oven at 85°C overnight.

### Fabrication and integration of TEER measurement system into the microdevices

The electrodes were designed using AUTOCAD program considering critical setup factors such as their size (150 µm) and the separation between them (150 µm) as well as the electrode-EC position to allow a homogeneous current density across the cell barrier. Gold electrodes were fabricated through photolithography technique on a glass slide. Initially, the coverslips were activated in the plasma chamber and then coated with the reversal photoresist AZ 5214 by spin coating. After that, the glass was soft-baked for 2 min at 95°C and exposed using an I-line mask aligner to transfer the design from the acetate mask. Then the resin was inversed by baked at 120°C for 2 minutes and immediately a flood exposure was carried out without the mask to fix the reverted angle. Finally, the resin was developed using the developer AZ 400K diluted in water in a ratio of 1:4. The metals deposition of Cr/Au 20/80nm over the exposed surface was performed by evaporation and the lift-off process was accomplished employing AZ 100 remover. Once fabricated the electrodes were cleaned with piranha solution for 10 minutes. PDMS replicas and the electrodes-glass slide were bonded together by plasma with precise alignment to place the electrode close to the endothelial barrier zone. Finally, the microsystem was left in the oven at 85°C overnight, obtaining the TEER measurement system integrated into the BBB-oC (TEER-BBB-oC).

### Cell seeding protocol

All steps, including cell culture, were performed in a sterile laminar flow hood, and only sterile materials, solutions and techniques were used. Human ECs were cultured in T75 culture flasks coated with collagen Type I from rat tail solution (25 μg/mL in PBS 1x) and supplied with endoGRO™ medium with 1 ng/mL of bFGF. Human pericytes and astrocytes were cultured in T75 culture flasks coated with poly-L-lysine (2 μg/cm^2^ in sterile water) in the corresponding pericyte and astrocyte media. All cells were kept in a humidified incubator at 37°C and 5% pCO_2_ and the cell media were changed every two days. Cells were detached using 0.25% trypsin/EDTA for EC and 0.05% trypsin/EDTA for pericytes and astrocytes. For further experiments, cells were used until passage 5 for astrocytes and until passage 7 for pericytes and EC.

To mimic the brain neuronal-vasculature network on the chip we adapted a procedure described by Campisi and collaborators [22]. As depicted in Scheme 1, pericytes and astrocytes embedded in a hydrogel were injected into the central chamber of the chip and after two days, ECs were introduced in one of the lateral channels. For this purpose, human pericytes and astrocytes were trypsinized and 40.000 of each cell type were mixed. This mixture was centrifuged at 1000 g for 5 min, and the pellet was maintained at 4°C while the fibrin hydrogel was prepared. For this purpose, 50 µL of filtered fibrinogen 3 mg/mL in PBS and 1 µL of thrombin 100 U in PBS were mixed with the cell pellet and injected into the central chamber. The microdevice was incubated for 15 min at 37°C and 5% pCO_2_ to allow the hydrogel polymerization and then, both channels were supplied with a mixture of endothelial and astrocytes medium in a 1:1 ratio (EM:AM medium). After two days, one of the lateral channels was coated with collagen (25 μg/mL in PBS 1x) for 50 min at 37°C and 5% pCO_2_ while 100.000 EC cells were detached and centrifuged at 1000 g for 5 minutes and resuspended in EM:AM medium. Then, the cells were injected in the collagen-coated channel and the chip was flipped for 1,5 h at 37°C and 5% pCO_2_ to allow the attachment of the cells to the hydrogel wall. Lastly, EM:AM medium was supplied on both channels and the microdevices were maintained in the incubator for 5 days before conducting any experiment. Cell culture medium was changed daily.

**Schema 1.**
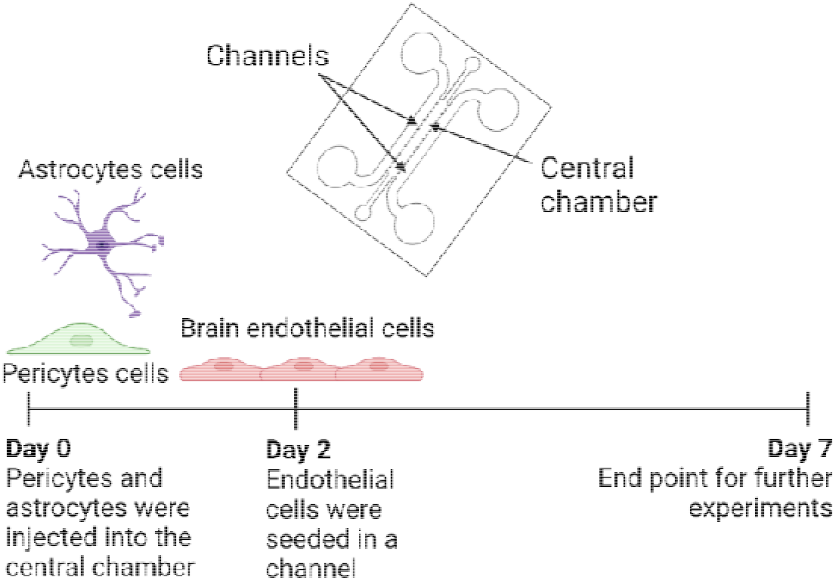
Timeline of BBB-oC cell seeding.

### Characterization of assembled BBB

#### Bright-field microscopy images

On day 7 of culture, optical images of the BBB-oC microdevice were taken using an inverted optical microscope (Olympus IX71) with an integrated CCD Hamamatsu camera. All the images were further analyzed and processed by ImageJ/Fiji ® software (NIH, USA)

#### Characterization by Fluorescent label

To establish the morphology and position of the neurovascular cells in the BBB-oC, F-actin was labelled with phalloidin-iFluor 594 and the cell nuclei with Hoechst 33342. For this purpose, cells in the BBB-oC on day 7 were washed with PBS 1x and fixed with commercial formalin solution (4% w/v formaldehyde) for 15 min at room temperature (RT). Fixed cells were rinsed three times with 1.5 mg/mL of glycine in PBS 1x (PBS-G) and later permeabilized with 0.1% TritonX-100 in PBS-G for 10 min at RT. Afterward, the cells were washed three times with 0.1% TritonX-100 in PBS 1x (PBS-T) and incubated with phalloidin-iFluor 594 (1:1000) in 1% bovine serum albumin (BSA) in PBS-T for 1.5 h at RT. Then, the cells were rinsed with PBS-T and incubated with Hoechst 33342 (1:1000) in PBS-T for 15 min at RT. Finally, the cells were washed three times with PBS 1x and the BBB-oC was observed by confocal microscopy (Leica TCS SP5 Multi-photon system) at 63x.

In addition, we evaluated the cell viability every day after the EC seeding through live/dead assay using calcein-AM to stain alive cells and ethidium homodimer-1 (EthD-1) for dead cells according to an adaptation of the manufacturer protocol. A mixture of calcein-AM 2 μM and EthD-1 4 μM in PBS 1x was prepared just before use (live/dead solution). Cells into the BBB-oC were washed with PBS 1x and live/dead solution was added and incubated for 1 h at RT. Then, the device was washed carefully with PBS 1x twice and finally filled with PBS 1x and the microdevices were visualized by fluorescent microscopy (Olympus IX71) at 10 and 20x. Fiji/ImageJ® software was then employed to analyze the acquired images.

We also assessed the cytotoxic effect of the GNR-PEG-Ang2/D1 over the cells in the BBB-oC microdevice using live/dead assay. On day 7, cells were rinsed with PBS 1x and incubated with GNR-PEG-Ang2/D1 0,4 nM or GNR covered with cetyltrimethylammonium bromide (GNR-CTAB) 0,1 nM that was previously reported as cytotoxic at this concentration [43] for 24 h at 37°C and 5% CO_2_. Controls were conducted using only EM:AM medium taking the images at timepoints 0 and 24 h. After 24 h, the live and dead assay was conducted, and the EC zone was observed at 10x.

#### Characterization with Immunofluorescence

After fixation and permeabilization as previously described for fluorescent labelling, cells were washed three times with PBS-T and blocked with 3% BSA in PBS-T for 1 h at RT. The cells were incubated with antibodies against TJs proteins such as ZO-1 (1:100) and VE-cadherin (1:100) in 3% BSA in PBS-T overnight at 4°C. After that, cells were washed three times with PBS 1x and incubated with the corresponding fluorophore-conjugated secondary antibody anti-rabbit Alexa 568 (1:1000) and anti-mouse Alexa 488 (1:1000) in 1% BSA in PBS-T for 2 h at RT. Then, the chip was washed with PBS-T and incubated with Hoechst 33342 (1:1000) for 15 min at RT. Finally, the cells were washed three times with PBS 1x and the microdevices were inspected by confocal microscopy at 63x (Leica TCS SP5 Multi-photon system). Fiji/ImageJ® software was then employed to treat the acquired images.

To determine if GNR-PEG-Ang2/D1 incubation has any influence over the endothelium, immunofluorescence of ZO-1 and VE-cadherin was performed. At day 7, the device was washed with PBS 1x and incubated with GNR-PEG-Ang2/D1 0,4 nM for 24 h at 37°C and 5% pCO_2_. Controls were conducted using GNR-CTAB 0,1 nM or with the equivalent quantity of the peptides Ang2 (175,6 nM) or D1 (69,2 nM) covering the surface of the GNR. After 24 h, ZO-1 and VE-cadherin were immunostained and imaged.

### Permeability assays

#### - Fluorescence imaging permeability assay

Optical permeability assays were carried out employing the fluorescence-tracers with different molecular weights sodium fluorescein (NaFI, 376 Da) and dextran70-FITC (D70, 70000 Da). In previous works, those standard tracers have been widely used to validate the permeability performance of the engineered BBB according to size-exclusion [19,21,40]. Fluorescent solutions of NaFI 100 µM and D70 100 µM were prepared in EM:AM medium to maintain the physiological conditions. On day 7 of BBB-oC culture, cells were washed with PBS 1x to remove cell debris and both channels were supplied with EM:AM medium to capture the time point 0 using the bright field and fluorescent settings of an inverted fluorescence microscope (Nikon Eclipse Ti2). Then, the corresponding fluorescent tracer was injected into the endothelial while EM:AM cell medium in the non-endothelial channel, and time-lapse images were captured every 3 min for 1 h. We used the microsystem without EC as a control to demonstrate that the permeability restriction is due to the assembled BBB. Fiji/ImageJ® software was used to analyze the acquired images and the permeability coefficients (P) (cm/s) were determined as detailed by Campisi and collaborators [22]:

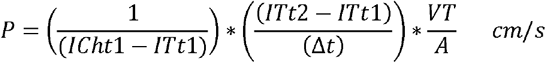

Where *ICht1 and ITt1* are the fluorescence intensity at the initial time in the endothelial channel and central chamber; respectively (u.a.), *ITt2* is intensity in the central chamber at end time (u.a.), Δt is the difference between the initial and the final time (s), VT is the volume of the central chamber (cm^3^) and A is the surface area of the endothelial wall (cm^2^).

Once assessed the permeability of the BBB-oC with standard tracers, the permeability of GNR-PEG-Ang2/D1 across the endothelial barrier was analyzed. For this purpose, GNR was fluorescently labelled by linking Alexa647 hydrazide with its carboxyl groups from SH-PEH-COOH. Permeability assays were conducted by injecting the GNR-PEG-Ang2/D1-647 or GNR-PEG-D1-A647 0.4 nM as control into the endothelial channel, while EM:AM medium was added to the other channel. To analyze GNR permeability, time-lapse images were captured every 3 min for 1 h. In addition, since Ang2 peptide facilitates the BBB crossing mostly through LRP1-mediated endocytosis, we employed dynasore as an endocytosis blocker to assess Ang2 influence on GNR-PEG-Ang2/D1 passage. [44]. For this purpose, on day 7 of BBB-oC, the cells were rinsed thoroughly with medium and then pre-incubated with dynasore 80 µM in cell medium for 30 min in the incubator. All the steps of the experiments were conducted in EM:AM serum-depleted medium to avoid that dynasore binds to the serum proteins and losing its activity. Since the dynasore stock was solubilized in DMSO, a control with DMSO 1% in cell medium was performed to evaluate the impact of the solvent on the endocytic inhibition. Afterward, the medium was discarded and the GNR-PEG-Ang2/D1-A647 0.4 nM in dynasore 80 µM or DMSO 1% in cell medium were injected into the endothelial channel of the BBB-oC, while the non-endothelial channel was fulfilled with dynasore 80 µM or DMSO 1% in cell medium. Finally, images were taken every 3 min for 1 h.

#### - TEER permeability assay

TEER values from EC culture were read out by impedance spectroscopy using the TEER-BBB-oC device. For this purpose, TEER measurements were conducted in sterile conditions inside the cell cabinet with the SP-150 Bio-Logic potentiostat instrument (Figure 1S, Supplementary). The impedance spectrum was recorded by connecting the working electrode in the endothelial channel and the counter/reference electrode connected in the central channel where the hydrogel was injected. Impedance spectra were recorded using an alternating current (AC) with a sinus amplitude of 25 mV ranging from 10 Hz to 1 MHz and 10 readings were registered for each measurement. Prior to TEER assessment, EM:AM media was refreshed in both channels and waited for 15 min before TEER measuring to stabilize the recording temperature [45]. TEER values were plotted as Nyquist and the equivalent circuit was modeled for fitting the data with Zview 4® software (Scribner, USA) to determine the changes in electrical resistance and capacitance due to the cell barrier over days.

TEER values were determined to assess the changes in electrical resistance due to the interaction between the endothelial barrier with GNR-PEG-Ang2/D1 0,4 nM for 24 h at 37°C and 5% pCO_2_. Controls were conducted using only EM:AM medium, GNR-PEG-CTAB (0,1 nM), Ang2 (175,6nM) or D1 (69,2nM) peptide in EM:AM cell medium to evaluate their influence over the TEER.

### Synthesis and characterization of GNR-PEG-Ang2/D1

The GNR-CTAB was functionalized with polyethylene glycol (PEG), Ang2 and D1 peptides. Briefly, GNR-CTAB was synthetized by a seed-mediated growth method [46,47]. The GNR-CTAB was purified by centrifugation at 16000g for 30 min and the pellet was resuspended in Milli-Q water. Then, GNR-CTAB was conjugated with two types of polyethylene glycol (PEG): HS-PEG-OMe and HS-PEG-COOH to stabilize it by charge and add terminal groups that allow further functionalization with Ang2 and D1 peptide. The adsorption of the PEGs onto the GNR-CTAB surface was performed following a previously described protocol [38,48]. We functionalized the GNR-PEG with the Ang2 and D1 peptides employing the terminal-carboxyl group from the PEG molecules with amine groups from the peptides through a coupling reaction with EDC/NHS [37]. Finally, D1 and Ang2 peptides were incubated overnight at RT and then the final product GNR-PEG-Ang2/D1 was purified to eliminate peptide-free molecules obtaining.

To characterize the obtained nanosystems, UV-visible absorption spectra were recorded to determine the surface plasmon band with multiplate reader (Infinite M200 PRO Multimode). The measurement was performed in the wavelength range of 400 to 900 nm at RT and Milli-Q water was used as blank. In addition, dynamic light scattering (DLS) was used to determine hydrodynamic diameter (Dh) and poly dispersion index (pdi), and surface charge (zeta potential) was measured by laser Doppler micro-electrophoresis both using ZetaSizer equipment (Nano ZS Malvern). Moreover, transmission electron microscopy was employed to determine the size and morphology of the nanoparticles. The samples were deposited on copper grids with a formvar/carbon and were observed employing electron microscopy (JEOL J1010) using an Orius CCD camera, and the particles were measured and counted by Fiji/ImageJ® software.

#### Cytotoxic assays of GNR-PEG-Ang2/D1

To determine non-cytotoxic concentration range of GNR-PEG-Ang2/D1 for all the cells conforming the BBB-oC (ECs, astrocytes and pericytes), flow cytometry using DAPI and Annexin V-Alexa647 was employed. Dead cells are marked with DAPI while cells at the early apoptosis stage are marked by Annexin V-Alexa647. For this purpose, cells were detached and 50.000 astrocytes or pericytes cells were seeded in poly-L-lysine (2 μg/cm^2^ in sterile water) or 50.000 EC in collagen (25 μg/mL in PBS 1x) coated 24-well plate with their corresponding medium and were maintained for 24 h at 37°C and 5% pCO_2_. Then, different concentrations of GNR-PEG-Ang2/D1 (0.05, 0.1, 0.2 and 0.4 nM) were added and left in the incubator for 24 h. Live and early apoptosis controls were performed with only EM:AM medium and staurosporine 1 µM, respectively. After that, cells were detached from the wells and were centrifuged at 1000 g for 5 minutes. The resulted pellet was resuspended in a solution with DAPI (1:2000) and anti-annexin V (1:2000). Finally, the samples were measured through flow cytometry with a cytometer equipment (GALIOS) and the results were analyzed by the Flowjo® software (Flowjo, USA).

### Statistical analyses

The data obtained from the cytotoxicity assays by flow cytometry, and permeability assays for the standard traces and GNR/endocytosis-blocking were analysed using One-way ANOVA with Dunnett’s and Tukey’s post-hoc test, respectively. On the other hand, the permeability assay of GNR-PEG-Ang2/D1 and GNR-PEG-D1 was analyzed using the unpaired statistical T-test. All the statistics were conducted with the Graphpad prism 8® software (GraphPad Software Inc, USA).

## Results and discussion

### Fabrication of microfluidic devices and integration of TEER system

To fabricate the BBB-oC, a compartmentalized device was designed inspired by the model developed by Farahat and collaborators [49]. As Figure 1A displays, the design is composed of a central chamber enclosed by trapezoidal posts with 100 µm between them, as well as two lateral channels to the central chamber. The posts can confine a hydrogel matrix in the central chamber without leaks due to the high hydrophobicity of the PDMS surface in relation to the hydrogel [50]. In addition, the two lateral channels operate as cell medium suppliers, or for endothelial culture in our case, and the reservoirs avoid media evaporation. The design was then engraved on a silicon wafer and replicated in PDMS. Finally, the bonding between PDMS replicas and the glass slides was done and we achieve multiple microdevices suitable for cell seeding and endothelial barrier assays (Figure 1D).

**Figure 1.**
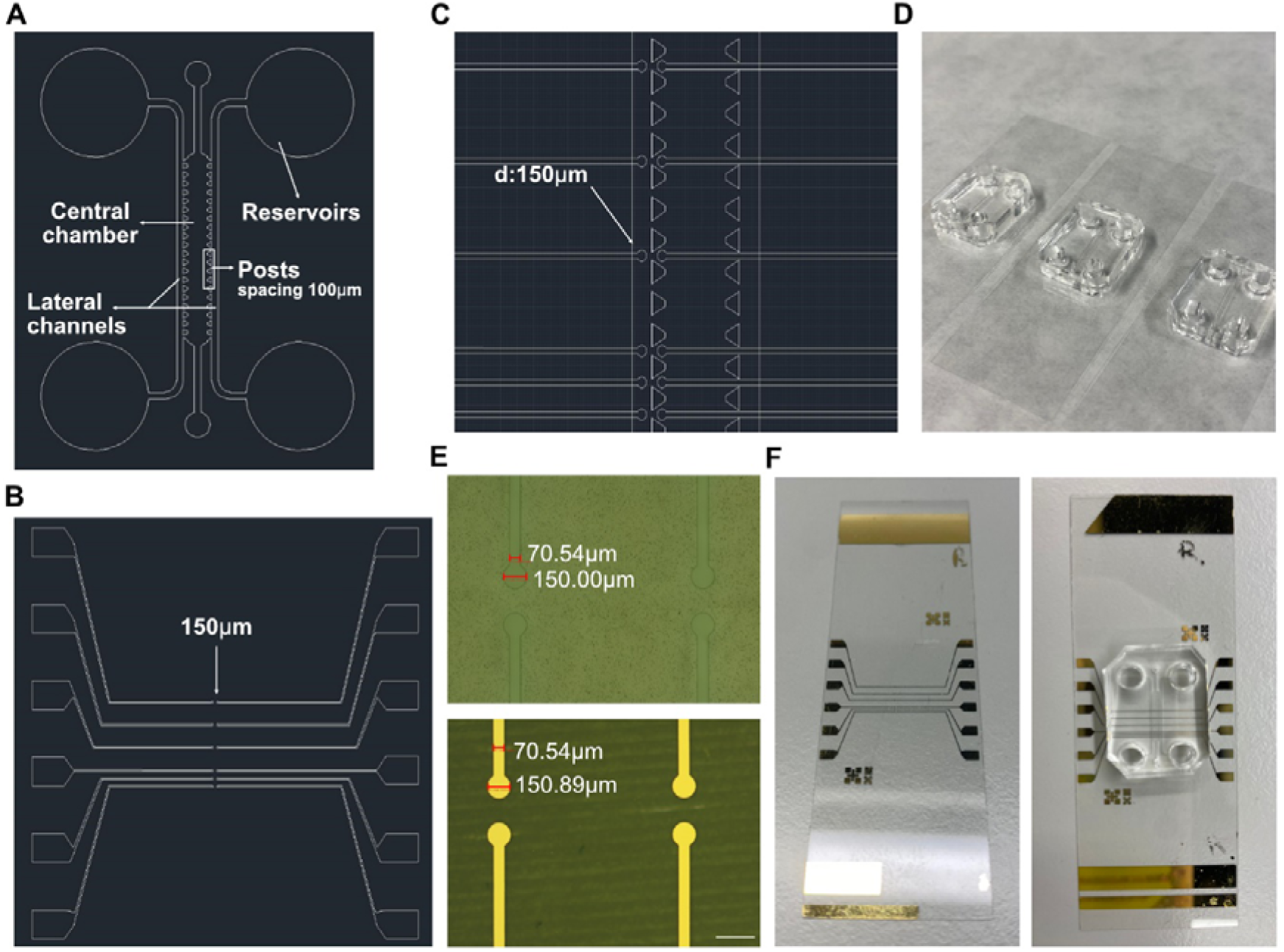
BBB-oC and measuring system design and fabrication. A), B), C) AutoCAD picture of the microfluidic design, electrodes design and its arrangement with the BBB-oC, respectively D) Fabricated BBB-oC devices suitable for cell seeding E) Optical images before and after Au/Cr evaporation on the lithographed design. Scale bar 250µm. and F) Finished TEER-BBB-oC ready for cell incorporation and resistance measuring.

TEER is a quantitative method for assessing the endothelial barrier permeability. Gold electrodes were designed considering multiple factors that could affect the correct TEER read-out [28]. Consequently, our electrode has a diameter and separation distance of 150 µm (Figure 1B) based on our endothelial zone (width of 100 µm) to reach a uniform current density on the barrier. In addition, the electrodes were positioned as closely as possible to the cell barrier (75 µm) which is in an equal distance between the electrodes (Figure 1C). Moreover, our design allows TEER measurement at multiple regions from the barrier on a single chip, providing information along the endothelial channel but also permits a perfect visualization of the EC barrier conversely to previous works which placed the electrodes over the cells tampering the imaging [18,19]. The gold electrodes were fabricated by photolithography; Cr/Au evaporation over the engraved design (Figure 1E) and a lift-off process were conducted resulting in a gold electrode with a height of 98 nm. Then, the electrodes were aligned-perpendicular to the endothelial zone of the PDMS and finally bonded by plasma treatment obtaining a successful TEER-BBB-oC system as can be seen in Figure 1F.

### Characterization of the neurovascular network into microfluidic devices

To mimic the brain neurovascular network, human pericytes and astrocytes cells were included in a fibrin hydrogel as ECM scaffold in the central chamber of the chip and human ECs in one of the lateral channels (Figure 2A). Fibrin is a natural hydrogel that can mimic a low stiff supporting extracellular matrix that is suitable for hosting vascular networks [51]. It is formed by the action of the protease thrombin, which cleavages the protein fibrinogen leading to fibrinogen monomers that self-associate into a stable fibrin mesh by the action of disulfide bridges with no need for radiation or additional crosslinking reagents [52]. After the tri-culture cell seeding, optical bright field images were taken at the endothelial and neuronal zone of the BBB-oC on day 7. ECs are accumulated on the hydrogel-channel interface of the device allowing close contact with the pericytes and astrocytes, as the natural BBB (Figure 2B), which are embedded in the 3D scaffold in the central chamber (Figure 2C). F-actin from the cytoskeleton in the BBB-oC was stained with phalloidin-iFluor 594 to confirm the cell morphology and position in the microdevice. Figure 3D shows the characteristic star-shape of astrocytes and pericytes cells embedded in the fibrin matrix and the EC located at the interface between the hydrogel and the channel.

**Figure 2.**
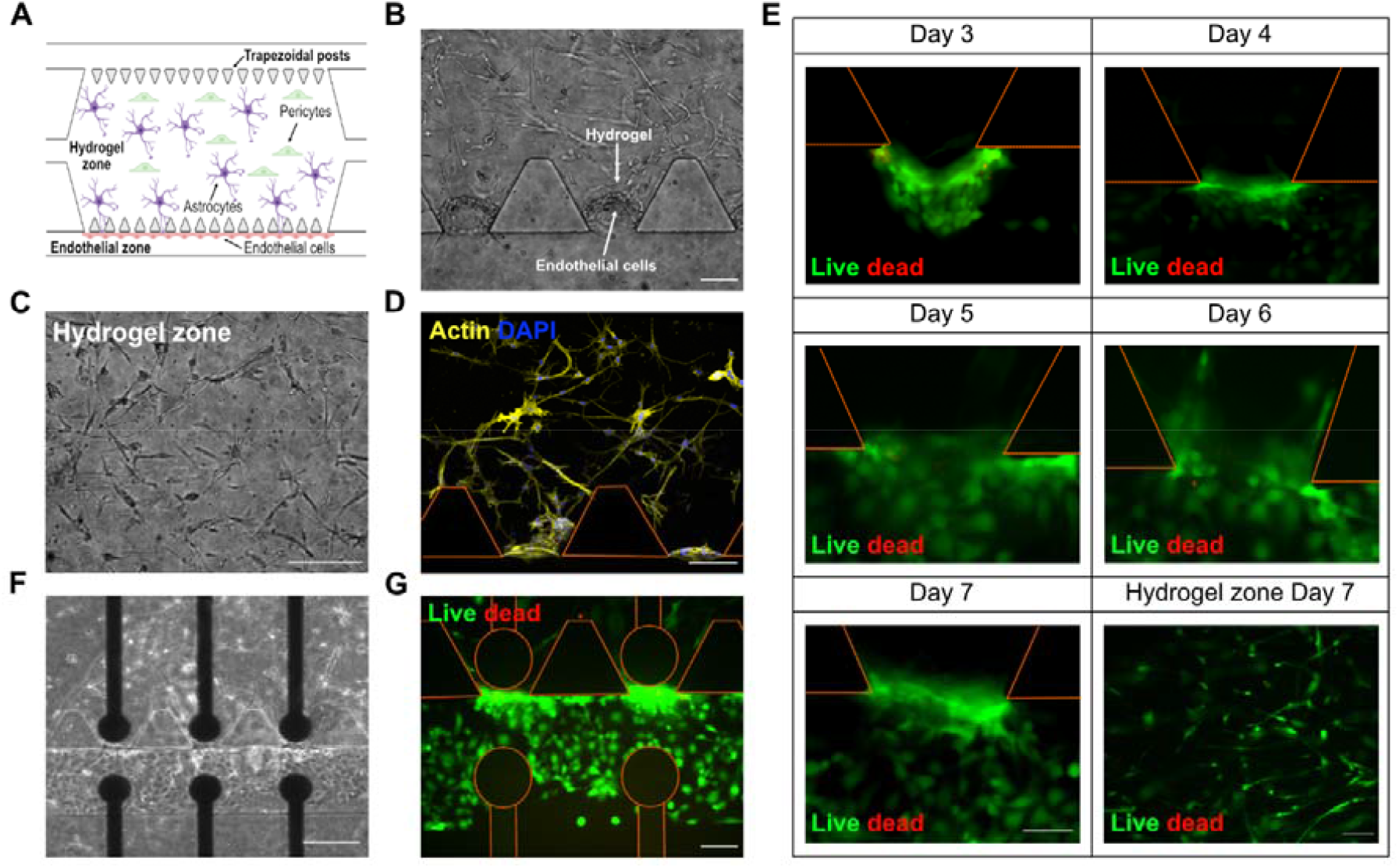
BBB-oC devices after cell seeding with tri-culture. A) Schematic representation of neurovascular cells arrangement into the BBB-oC B) Optical images at Day 7 of ECs at 10x and C) Hydrogel zone at 20x D) Confocal z-projections of F-actin from cytoskeleton of neurovascular cells E) Cell viability assay of ECs every day from day 3 of cell seeding at 20x and hydrogel zone at Day 7 at 10x F) Optical images and D) cell viability assay of endothelial cells at Day 7 into TEER-BBB-oC after TEER measurement every day at 10x. Scale bar 100μm excepting E) scale bar 50μm.

**Figure 3.**
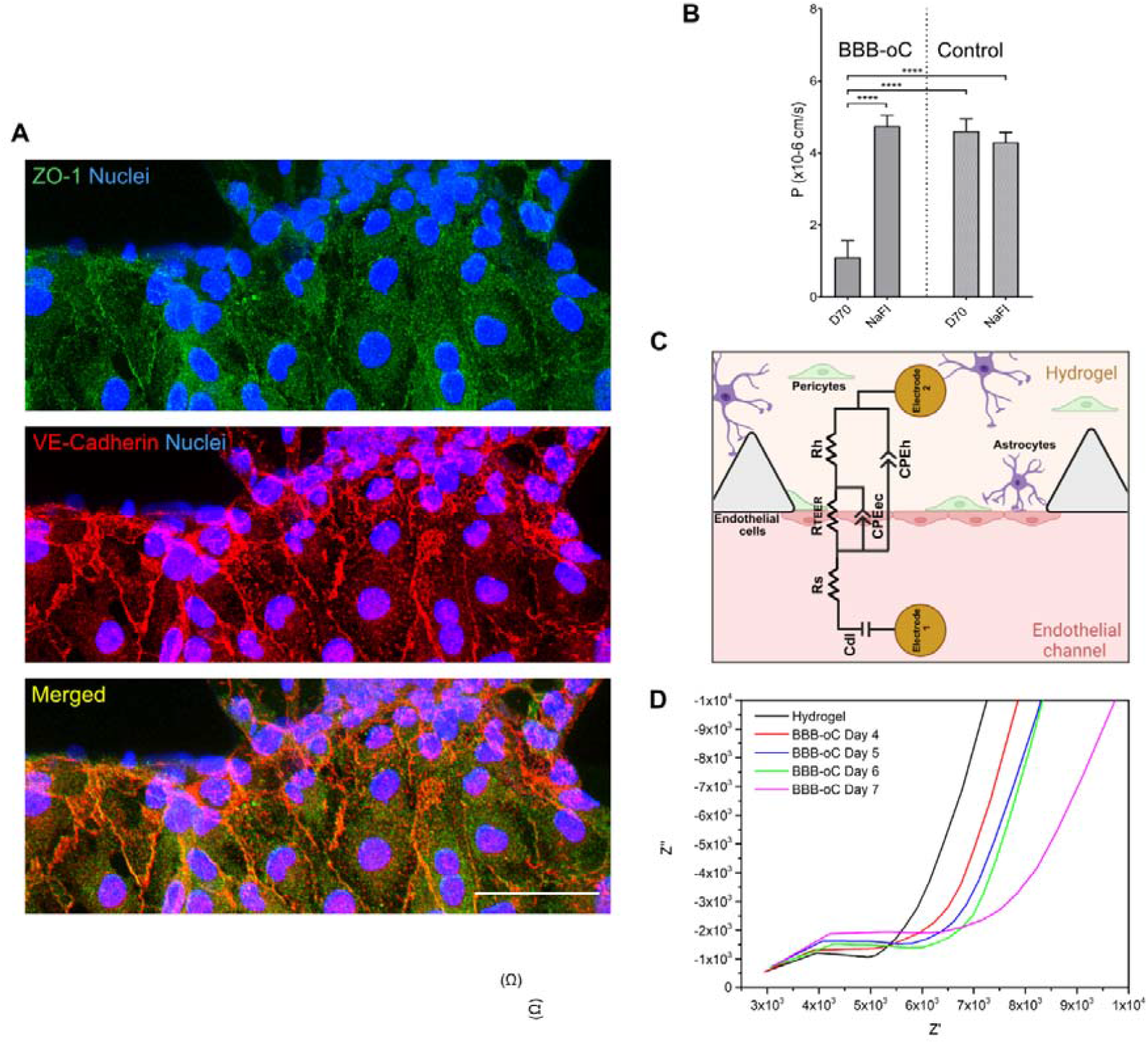
BBB-oC characterization. A) Confocal z-projections of ZO-1 and VE-cadherin between endothelial cells into BBB-oC devices. Scale bar 50 μm. B) Permeability assays of fluorescent standard molecules with different molecular sizes into the BBB-oC and control without endothelial barrier (N=3) **** p <0.0001 C) Equivalent circuit diagram that can be applied to our BBB-oC D) Nyquist plot of TEER measurement of BBB-oC before (only hydrogel with astrocytes and pericytes) and after endothelial cell seeding at Day 4, 5, 6 and 7.

On the other hand, live/dead assays were performed to evaluate the viability of the BBB-oC over time. As shown in Figure 2E, the green signal from the ECs visibly indicates that the cells are alive nearly at 100% throughout the time until day 7 as well in the hydrogel zone most of the cells are viable. Therefore, the use of fibrin as ECM for fragile cultures such as human astrocytes and pericytes and ECs, considering the culture conditions used, allow to have fully viable cells in the BBB-oC to perform any assay. TJs structures linking the ECs were formed inside the chip. Indeed, to evaluate the BBB development and the appropriate formation of the TJs between the ECs, ZO-1 and VE-cadherin were immunostained and observed under a confocal microscope at the BBB zone. Zonula occludens 1 (ZO-1) is a scaffolding protein that connects the TJs proteins between actin from the cytoskeleton structure in cell-to-cell interactions [53]. On the other hand, VE-cadherin is a strictly endothelial-specific adhesion molecule located at junctions between endothelial cells [54]. Figure 3A markedly evidenced the presence of ZO-1 stained in green and VE-cadherin in red between adjacent endothelial as well as the merged.

We carried out permeability assays using fluorescent tracers with different molecular weights. On day 7, NaFI (376Da) or D70 (70000 Da) were injected into the endothelial channel and fluorescent images at the neuronal area were recorded. As shown in Figure 3B the permeability values obtained from the BBB-oC suggest a size-dependent exclusion, where smaller molecules go faster through the BBB than the bigger ones. In the platform without ECs (control) the obtained coefficients show significant higher values of permeability for both NaFI (4,30 ×10^−06^ cm/s) and D70 (4,61 ×10^−06^ cm/s) in comparison with the tri-cultured cell system for D70 (1,11 ×10^−06^ cm/s) but not for small molecules as NaFI (4,76 ×10^−06^ cm/s). Therefore, these molecules are almost equally fast towards the neuronal zone when our constructed barrier is absent, thus shows no size restriction capability. The permeability values obtained with our developed BBB-oC are in the order of 10^−6^, which agrees with previous values reported in the literature for microfabricated 3D models [55], despite our system exhibited higher restriction against molecules as dextran than previous works that used the same EC line [56,57].

The permeability of the constructed EC barrier was also analyzed by TEER measurements. For impedance spectroscopy measurements, an AC voltage at swept frequency is applied between two electrodes on each side of the BBB distanced by 150 µm, very close to the barrier to reduce the total resistance of the system. With this technique is measured the amplitude and phase of the resulting AC current after crossing the BBB barrier. To analyze quantitatively the results obtained with TEER, the impedance measurements were fitted in an electrical circuit that defines the behavior of the measured platform. As Figure 3C shows, we considered for the electrical circuit the resistance (R_teer_) produced by the TJs between EC (paracellular) but also the contribution from EC layer (transcellular) described in our system as a non-ideal capacitance (CPE_ec_) which is frequency dependent. In our platform, we also considered other constant factors in the circuit, such as the double layer capacitance generated on the electrodes surface (C_dl_) and the resistance of the culture media solution (R_s_), but most importantly, the resistance and capacitance generated by the hydrogel with astrocytes and pericytes (R_h_ and CPE_h_) [58]. Supplementary Figure S2 shows the circuits used and the fitting of the TEER results in the absence and presence of the BBB barrier with a fitting χ^2^ error of 2.10^−3^. This study was conducted every day during the formation of the barrier to monitoring the development of the TJs in BBB. Higher TEER values indicate that TJs between ECs are strengthened and thus permeability to the brain is reduced. The impedance values before the endothelial seeding (hydrogel with astrocytes and pericytes) show lower resistance and the total absence of the characteristic resistive behavior which arises and increases after the endothelial incorporation (Figure 3D). The Nyquist plots of the impedance log after the EC seeding on days 4, 5, 6 and 7 show the shift of the impedance curve towards higher Z’ values demonstrating a gradual increase in TEER resistance during barrier formation (Figure 3D). TEER values from 4500 Ω reached 10.400 Ω since day 4 to day 10, respectively. Previous works also have observed TEER increasing over days related to the TJs formation between ECs [21,59,60]. As previously commented, one of the advantages of our electrode design is the possibility to observe the cell culture with microscopy in the chip contrasted to other TEER setups. Figure 2F displayed the presence of the neurovascular network but also Figure 2G taken on day 7 exhibits the not cytotoxic effect due to the daily TEER measurements conducted over the cell culture.

### Characterization of GNR-PEG-Ang2/D1

First, we performed a synthesis of GNR covered with CTAB as a stabilizer by seed-mediated growth method. Then, GNR-CTAB was conjugated with PEG obtaining GNR-PEG which confers several advantages such as stability, biocompatibility, increase blood circulation time, and others [61–63]. Finally, peptides were conjugated to the GNR-PEG where D1 peptide acts as Aβ sheet breaker and Ang2 favors the shuttling across the BBB mainly by LRP1 receptor (GNR-PEG-Ang2/D1). As Supplementary Figure S3A depicted, the UV-vis-NIR spectra of GNR-CTAB, GNR-PEG and GNR-PEG-Ang2/D1 revealed two characteristic plasmonic bands of absorption at around 510nm (longitudinal) and 740nm (transversal) as have been observed before in the literature [64]. In addition, the change in the Dh and zeta potential values confirmed the functionalization of the GNR-CTAB surface (Supplementary Figure S3B). On the longitudinal side, the Dh changed from 55nm to 74nm due to the PEG adsorption onto the surface through the thiol-Au bond but then it was modified to 60 nm after peptide functionalization. This behavior can be explained by the flexible character of PEG chains which can be contracted by peptide conjugation following an EDC/NHS protocol. Dh from the transversal side was also changed from about 1 to 4 nm. On the other hand, the electrokinetic surface potential (Z Potential) fluctuated from positive to negative charge (+37 to -8 mV) indicating that the nanoparticles were effectively functionalized. Firstly, GNR-CTAB displayed a positive potential as the cationic surfactant molecule CTAB is on the surface. This highly positive value (+37 mV) confers good stability to the nanoparticles due to the electrical repulsion between them. Then, CTAB molecules were replaced by PEG chains which have terminal groups negatively charged (methyl and carboxyl) at pH 7.4, thus conferring negative values (-32 mV) on the surface and providing good stability by electrical repulsion. Finally, the functionalization with Ang2 and D1 showed values near neutrality (-8 mV) due to the functional groups present in each peptide that neutralize the previous charge. This shift certainly decreases the electrical repulsion between the nanoparticles but gives relevance to steric stabilization by the molecules attached to the GNR surface. Finally, electron microscopy displayed the shape and size of GNR-CTAB, GNR-PEG and GNR-PEG-Ang2/D1 exhibiting a length of 35.7±3 nm and width of 10.5±1 nm (Supplementary Figure S3C). The analyzed images revealed a histogram of the distribution of their aspect ratio between the longitudinal and the transversal side of the GNRs which is mainly 3 (Supplementary Figure S3D).

### Cytotoxicity assays of GNR-PEG-Ang2/D1

The cytotoxicity of GNR-PEG-Ang2/D1 was evaluated in plate-cultured cells at different concentrations in EC, pericytes and astrocytes cells for 24 hours to determine a safety concentration range to administer into the microdevices. For this purpose, we used flow cytometer labelling with DAPI for the dead cells and Annexin-V for cells at the early apoptotic stage. Also, live and early apoptosis controls were assessed with cell medium and staurosporine 1µM, respectively. As Figure 4 exhibited, GNR-PEG-Ang2/D1 did not show any significant cytotoxic effect in EC, pericytes and astrocytes cells for the given range of concentration when comparing the % of cell viability with the live control. These results provide a cell safety range between 0,05 – 0,4 nM to apply in the microdevice.

**Figure 4.**
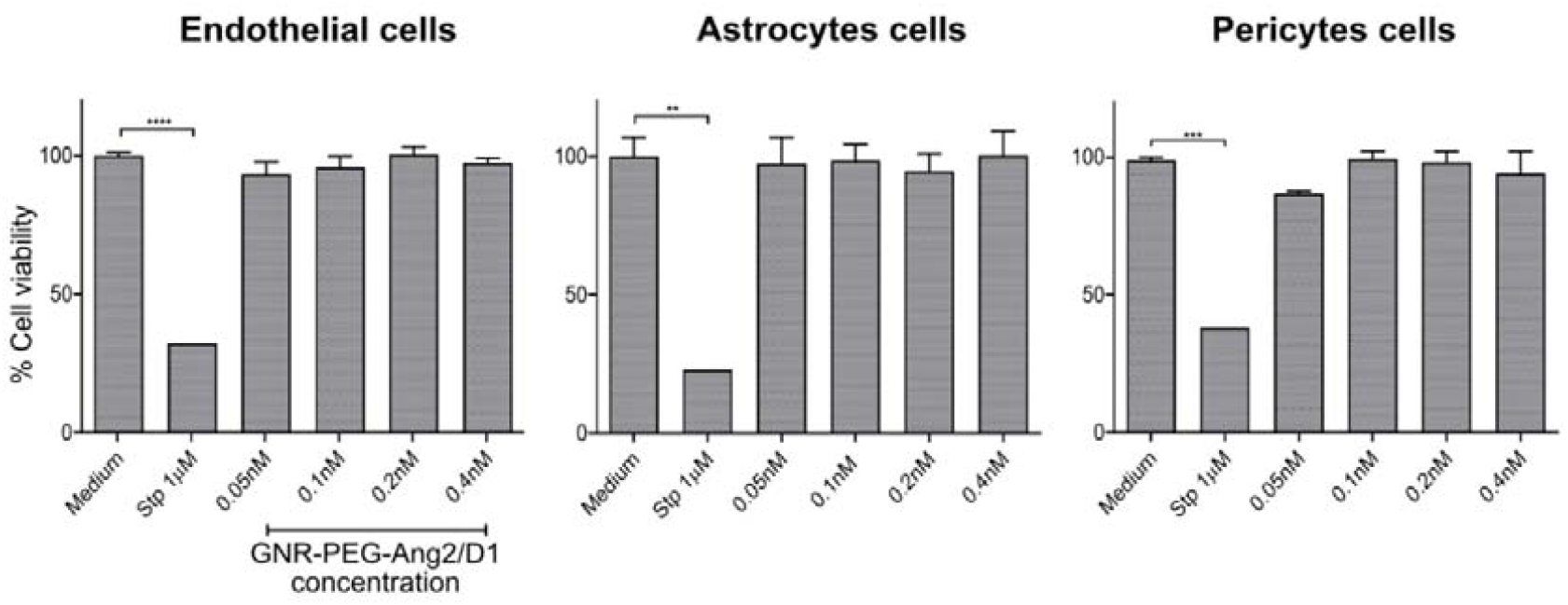
Cell viability assay of EC, astrocytes and pericytes cells incubated with GNR-PEG-Ang2/D1 at different concentrations for 24 h. Live control corresponds to the cells with only their culture medium and the death control corresponds to cells incubated with staurosporine 1 µM (N=3) **** p <0.0001, ***p <0.001 and **p <0.01.

Furthermore, we assessed the cytotoxic effect of GNR-PEG-Ang2/D1 on the endothelial barrier in the BBB-oC using a live/dead assay. On day 7, cells were incubated with GNR-PEG-Ang2/D1 0.4 nM for 24 h, based on the results performed in 2D cell cultures. Also, controls were conducted using GNR-CTAB 0,1 nM as a cytotoxic agent, and only EM:AM medium at timepoints 0 and 24 hours. Figure 5 shows a mainly green signal indicating that cells are alive after 24 hours in contact with GNR-PEG-Ang2/D1 0.4 nM, whereas GNR-CTAB cell death signal is visible, which agrees with previous data about the cytotoxicity of GNR-CTAB at this concentration [43]. Controls at t0 and t24h demonstrated that ECs in the BBB-oC do not die naturally due to extended culture times in the chip. Therefore, the interaction between the nanotherapeutic agent and the endothelial barrier displays fully viable cells in the BBB-oC.

**Figure 5.**
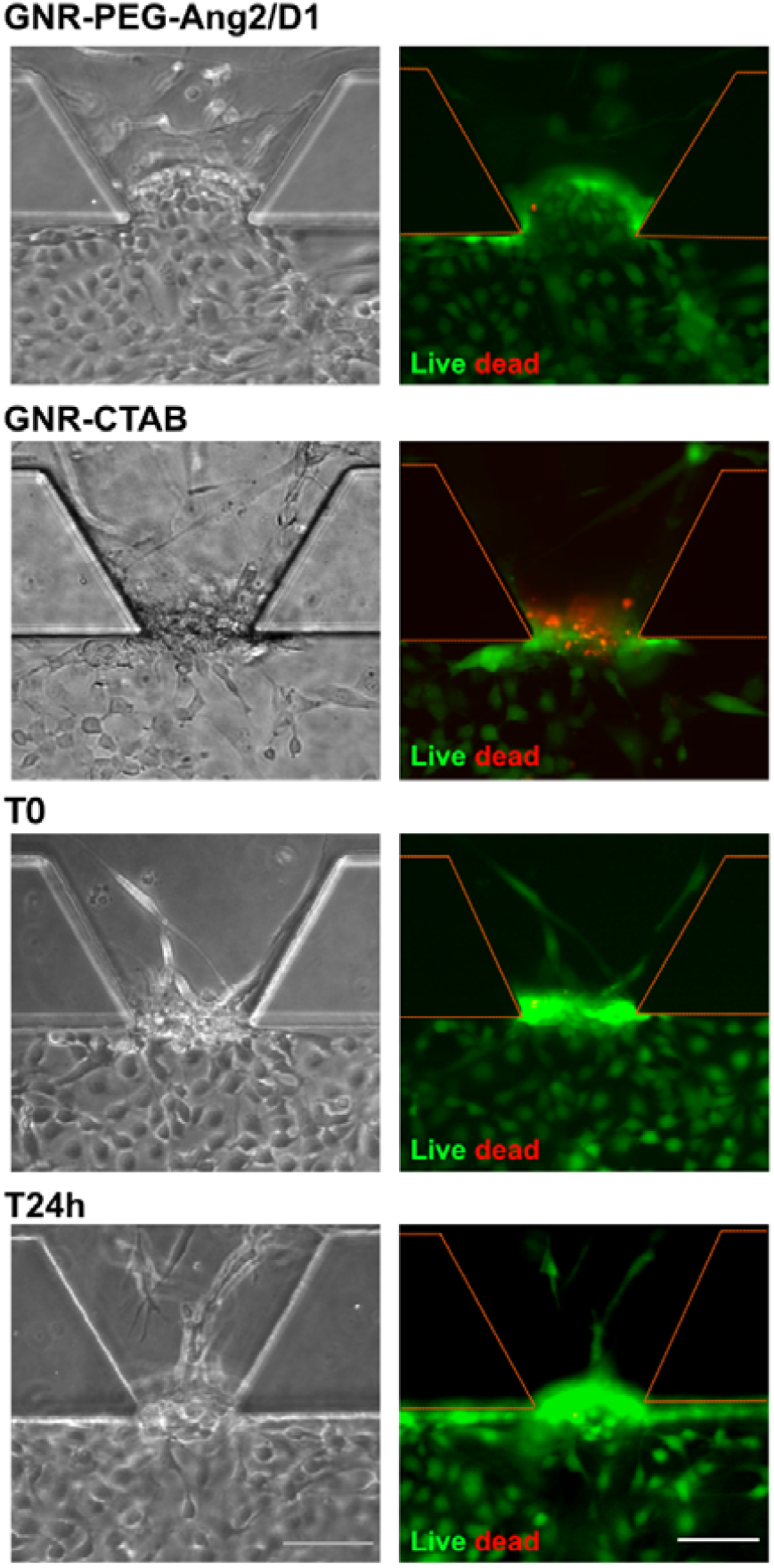
Cytotoxic evaluation into BBB-oC. Cell viability of EC was evaluated by incubating GNR-PEG-Ang2/D1 0,4nM for 24 h. Bright-field and fluorescent images were taken at 10x. Death control was achieved by exposing GNR-CTAB 0,1 nM for 24 h and also controls at time 0 and T24h. Scale bar 100 µm.

### GNR-PEG-Ang2/D1 permeability performance

To evaluate the ability of GNR-PEG-Ang2/D1 to cross the BBB, it was functionalized with the fluorophore Alexa647 and injected into the endothelial channel of the BBB-oC and fluorescent images were taken. Since Ang2 can penetrate the BBB by receptor-mediated endocytosis due to high LRP1 binding affinity, the expression of LRP1 in EC was confirmed by western blot [63] (Supplementary Figure S4) [65]. To evaluate the role of Ang2 in the entrance, nanoparticles without the shuttling peptide (GNR-PEG-D1) were used as control. Figure 6A reveals that after 1 h of incubation in the EC channel, the GNR-PEG-Ang2/D1 has a higher fluorescent signal in the neuronal area than GNR-PEG-D1. Also, as Figure 6B displays, permeability coefficients show that GNR-PEG-Ang2/D1 (4,74 ×10^−6^ cm/s) goes faster than GNR-PEG-D1 (3,02 ×10^−6^ cm/s) towards the neuronal zone. Finally, dynasore 80 µM dissolved in DMSO (0.2% DMSO) was added to block endocytic processes to prove the mechanism involved in the increased transport of the GNR-PEG-Ang2/D1. Permeability values (3,60 ×10^−6^ cm/s) displayed the reduction of the nanoparticle entrance when endocytosis was inhibited by dynasore (Figure 6C). Considering that the endocytosis inhibition leads to a decrease of the nanoparticles entry, it can be deduced that this mechanism is involved in the BBB crossing, but it should be noted that the cellular uptake pathways of nanoparticles depend on multiple factors such as size, surface charge, shape, among others, therefore the internalization of GNR-PEG-Ang2/D1 is not defined by this pathway alone [66–68]. Finally, DMSO control exposed that the GNR-PEG-Ang2/D1 transport is not affected by the dynasore solvent.

**Figure 6.**
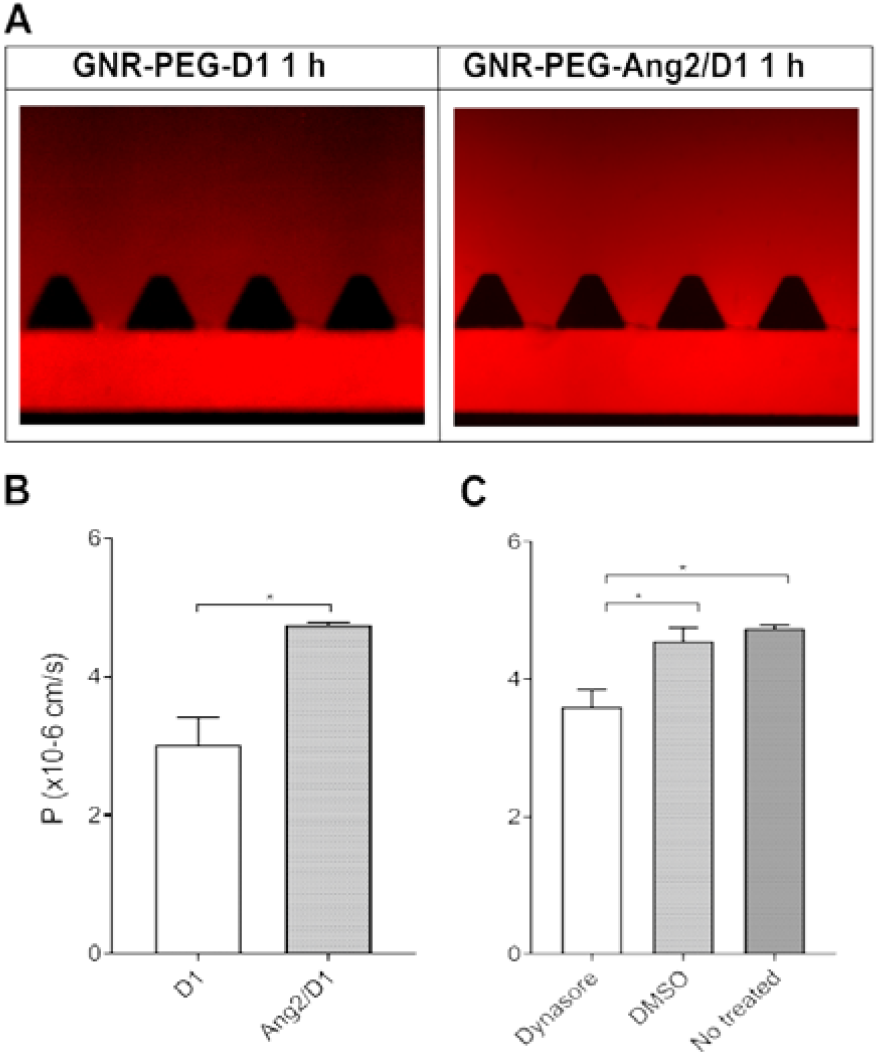
Permeability assays of GNR-PEG-Ang2/D1 into the BBB-oC. A) Fluorescent images after 1 hour of incubation in the EC channel B) P values of GNR-PEG-D1 and GNR-PEG-Ang2/D1 assessed for 1 h in BBB-oC (N=3) (*p <0.05) and C) P values of GNR-PEG-Ang2/D1 exposed to endocytosis inhibition (N=3) (*p <0.05). Controls were conducted with DMSO and cell medium as vehicles and no treated control, respectively.

After the incubation of ECs barrier with GNR-PEG-Ang2/D1, ZO-1 and VE-cadherin were immunostained and imaged by a confocal microscope. Controls were performed with only EM:AM cell medium and GNR-CTAB 0,1 nM for 24 h. Figure 7 exposed that ECs incubated with GNR-PEG-Ang2/D1 presents a higher fluorescent signal of ZO-1 and VE-cadherin compared with EM:AM control. Conversely, GNR-CTAB displayed lower VE-cadherin signal expression and practically no ZO-1 signal, mainly related to the cytotoxic effect of the nanoparticles at this concentration. In addition, to corroborate these obtained results, impedance recording was conducted before (Day 7) and after GNR-PEG-Ang2/D1 and controls incubation (Day 8). As Figure 7D depicted, Nyquist plot after the GNR-PEG-Ang2/D1 injection exhibited the shift of the curve to the right Z’ values, from 10.400 Ω on day 7 to 11960 Ω on day 8, indicating higher TEER values, which suggests greater formation of TJs and therefore more permeability restriction. Control with only EM:AM medium at day 8 revealed practically no differences in resistance due to longer culture time (Figure 7E). The administration of the cytotoxic GNR-CTAB 0,1 nM exhibited lower electrical resistance comparable with values before ECs seeding (Figure 7F) attributable to the cell membrane disruption produced by CTAB. Therefore, these results suggest consistency of the results observed in the immunofluorescence images and the TEER measurements and that both techniques work correctly. Furthermore, the results show a decrease in BBB permeability after GNR-PEG-Ang2/D1 crossover.

**Figure 7.**
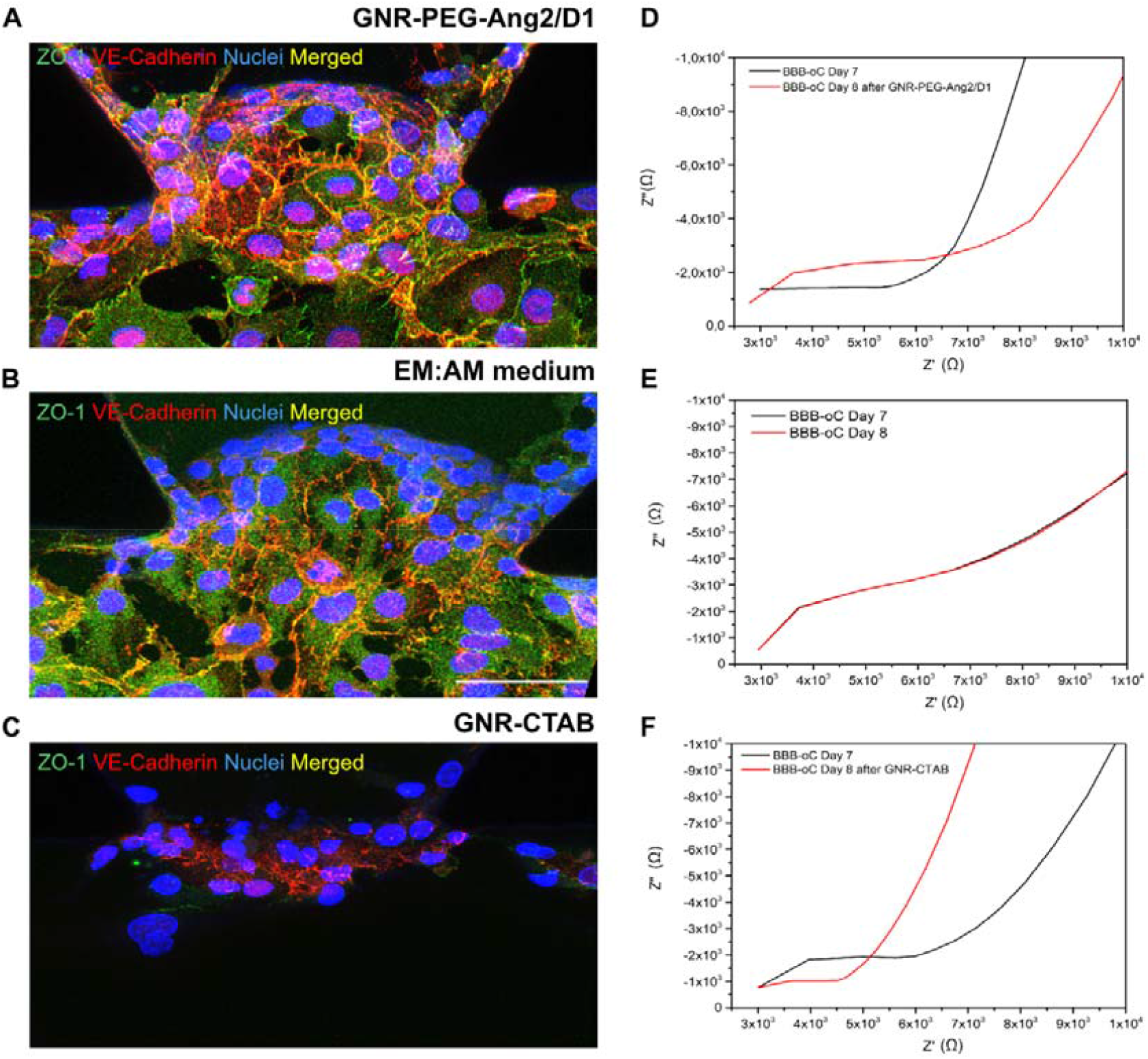
TJs influence after GNR incubation in BBB-oC. Confocal z-projections of ZO-1 and VE-cadherin between EC into BBB-oC devices on Day 8 after incubation with A) GNR-PEG-Ang2/D1, B) EM:AM medium only and C) GNR-CTAB for 24 hours (N=3). Scale bar 50 μm. Nyquist plot of TEER values of BBB-oC before (Day 7) and after 24 h of incubation of D) GNR-PEG-Ang2/D1, E) EM:AM medium only and F) GNR-CTAB.

To elucidate the components involved in the signal increase of TJs markers in the BBB after GNR-PEG-Ang2/D1 injection, controls were performed with only Ang2 or D1 peptides. Confocal images exhibited (Supplementary Figure S5) that D1 peptide produces an increment in the fluorescent signal of ZO-1 and VE-cadherin while Ang2 mainly displayed an increase in VE-cadherin signal, but not ZO-1. In agreement, Supplementary Figure S5 also shows an increase of electrical resistance when D1 peptide is administered but a slight decrease with Ang2. This reinforcement of TJs, which we observed in these results, has been previously reported with certain molecules such as hydrocortisone 550nM and cAMP 250µM, which are currently used for the development of suitable BBB models to achieve greater tightness between ECs [69,70]. But this behaviour has also been observed in peptides, as D1, but in different types of cells. The AMP77-97 peptide has been patented as a possible treatment for the oral mucositis, common in chemotherapy patients, due to its effect in stimulating the formation of new TJs between cells to repair the mucosal barrier [71]. In the case of NDDs, molecules that allow to reinforcement the TJs could give us possible attractive therapies considering circumstances such as AD, where there is a downregulation of the ZO-1 and occludins expression, as well as alteration in the assembly of claudin-5, leading to a BBB disruption and less brain protection, which means acceleration of disease progression [72–74]. Moreover, the possibility of studying the impact of nanoparticles on the brain endothelium in 3D in vitro models is interesting for the field of nanosafety and nanotoxicology due to antecedents that show alterations on the viability and integrity of claudin-5 and occludins in ECs by aluminium oxide nanoparticles [75].

## Conclusions

OoC technology is playing an essential role in reducing the number of animal experiments and offers suitable models needed to optimize the design of drugs, nanoparticles, and biopharmaceuticals products in a high throughput manner. In this direction, we have fabricated and tested a BBB-oC microdevice with an electrical resistance permeability sensor suitable for 3D co-culture analysis. We optimized a cell seeding protocol to incorporate human cells onto de chip to mimic the neurovascular network of the brain. Microscopy images showed the position, morphology, and full viability of the cells over days. In addition, immunofluorescence images displayed the development of TJs between adjacent EC, which are crucial for the low permeability of the BBB. Permeability assays revealed size exclusion ability and coefficient order according to the previously reported for microfabricated 3D models, indeed our system showed greater restriction compared with other works that used the same type of EC. TEER measurements showed an increase in electrical resistance over days until day 7. With this successfully developed BBB-oC platform, the permeability of a nanotherapeutic agent with potential for the treatment of AD was tested. The developed nanopharmaceutical is based on multi-decorated GNRs with the Ang2 peptide to encourage their pass through the BBB and D1 peptide to inhibit the Aβ fibrillation which is the main hallmark of AD. The GNR-PEG-Ang2/D1 was successfully synthesized and functionalized and the AD nanotherapeutic agent showed a non-toxic effect for the tri-culture at the given range of concentration (0,05-0,4 nM) for 24 hours. Live/dead assays into the BBB-oC determined that GNR-PEG-Ang2/D1 0,4nM for 24 hours is not cytotoxic for the 3D neurovascular system. Moreover, GNRs permeability assays revealed that Ang2 enhances the permeability of the GNR-PEG-Ang2/D1, and when endocytosis was inhibited, this mechanism was verified as responsible for entry. Finally, confocal images displayed increased ZO-1 and VE-cadherin signals as well as higher TEER values posterior to the GNR-PEG-Ang2/D1 injection into the BBB-oC. According to the results, D1 peptide seems to be involved and further experiments are needed to deeply understand this behavior. From these data, our TEER-BBB-oC system could be a useful tool to assess the permeability of new drugs more quickly and cheaper than in vivo models, but also could allow us to explore the influence of the nanotherapeutic agents over the CNS, opening new windows of amelioration in NDDs and accelerating the drug screening process.

## Acknowlegment

This work was supported by Networking Biomedical Research Center (CIBER), Spain. CIBER is an initiative funded by the VI National R&D&i Plan 2008–2011, Iniciativa Ingenio 2010, Consolider Program, CIBER Actions, and the Instituto de Salud Carlos III (RD16/0006/0012), with the support of the European Regional Development Fund. This work was funded by the CERCA Programme and by the Commission for Universities and Research of the Department of Innovation, Universities, and Enterprise of the Generalitat de Catalunya (2017 SGR 1079). This work was funded by the Spanish Ministry of Economy and Competitiveness (MINECO) through the project NEUR-ON-A-CHIP (RTI2018-097038-B-C21 and RTI2018-097038-B-C22). S. P. acknowledges support from BECAS CHILE. Also, thanks to MicroFabSpace and Microscopy Characterization Facility, Unit 7 of ICTS “NANBIOSIS” from CIBER-BBN at IBEC. As well, we would like to acknowledge Clara Alcon for her technical assistance in the performance of these experiments. MK acknowledge to Fondap 15130011 and Fondecyt 1211482. The schemes presented in this article were created with BioRender.com.

