## Supplementary material for "BBB-on-a-chip with Integrated micro-TEER for permeability evaluation of multi-functionalized gold nanorods against Alzheimer’s disease": Supplemetary Figures

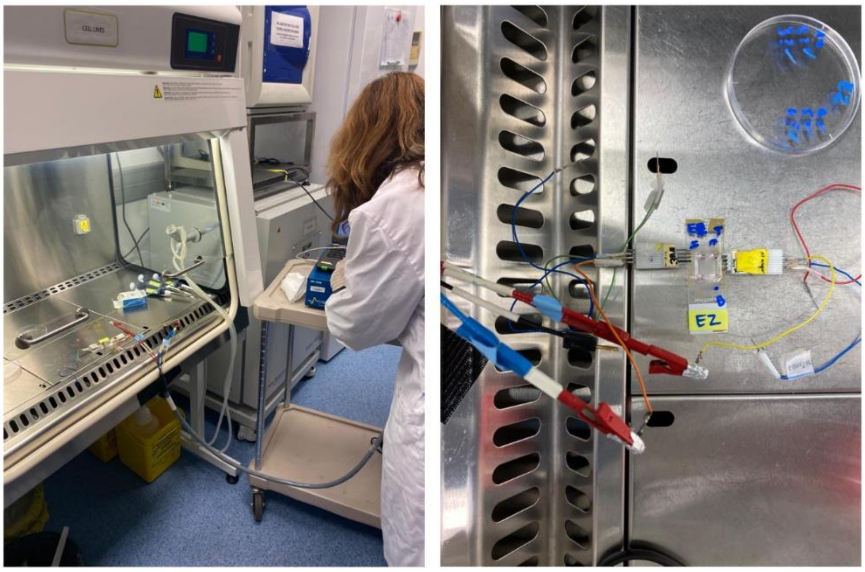


**Supplementary Figure S1.** Setup for TEER measurement in sterile conditions.

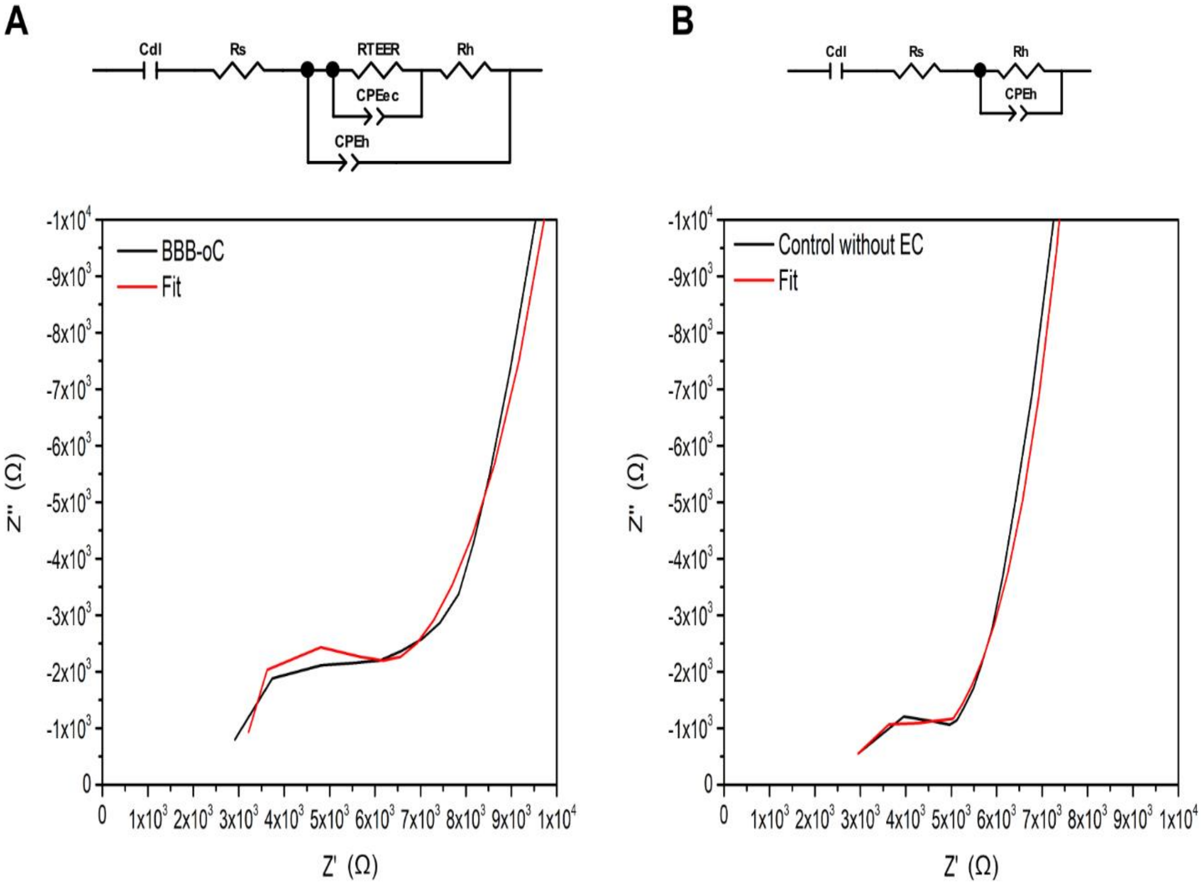


**Supplementary Figure S2.** Equivalent circuit and Nyquist plot of TEER from BBB-oC and control without EC and their fitting.

**Synthesis and characterization of GNR-PEG-Ang2/D1**

The GNR was functionalized with polyethylene glycol (PEG), Ang2 and D1 peptide, as previously described (Scheme 2). Briefly, GNRs covered with cetyltrimethylammonium bromide (GNR-CTAB) were synthesized by a seed-mediated growth method [55,56]. A seed solution was created by the reduction of $HAuCl_4$ 29.4 mM using $NaBH_4$ as a reducing agent in the presence of CTAB 0,1 M at 27°C in stirring conditions. Then, the 120 µL of seeds were added to a growth solution consisting of $HAuCl_4$, L-ascorbic acid and $AgNO_3$ in a matrix of CTAB 0.1 M and left them growing for 30 min at 27°C obtaining a red-brownish suspension. The GNR-CTAB was purified by centrifugation at 16000g for 30 min and the pellet was resuspended in Milli-Q water. Then, GNR-CTAB was conjugated with two types of polyethylene glycol (PEG): HS-PEG-OMe and HS-PEG-COOH to stabilize it by charge and add terminal groups that allow further functionalization with Ang2 and D1 peptide. The adsorption of the PEGs onto the GNR-CTAB surface was performed following a previously described protocol [57,58]. First, HS-PEG-OMe was added to GNR-CTAB solution at pH 12, to improve the Au-S bond, and was stirred for 10 min at RT. Then, the mixture was purified, and the pellet was diluted and adjusted at pH 12. Then, HS-PEG-COOH 1 mM was added and stirred for 1 h at RT and the resulted GNR-PEG solution was purified again. We functionalized the GNR-PEG with the Ang2 and D1 peptides employing the terminal-carboxyl group from the PEG molecules with amine groups from the peptides through a coupling reaction with EDC/NHS [44]. GNR-PEG was resuspended with a mixture of EDC and NHS in MES buffer at pH 5.5 for 15 min and [44] the mixture was purified. Finally, the pellet was incubated with D1 and Ang2 peptides in PBS buffer overnight at RT and then the final product GNR-PEG-Ang2/D1 was purified to eliminate peptide-free molecules obtaining.

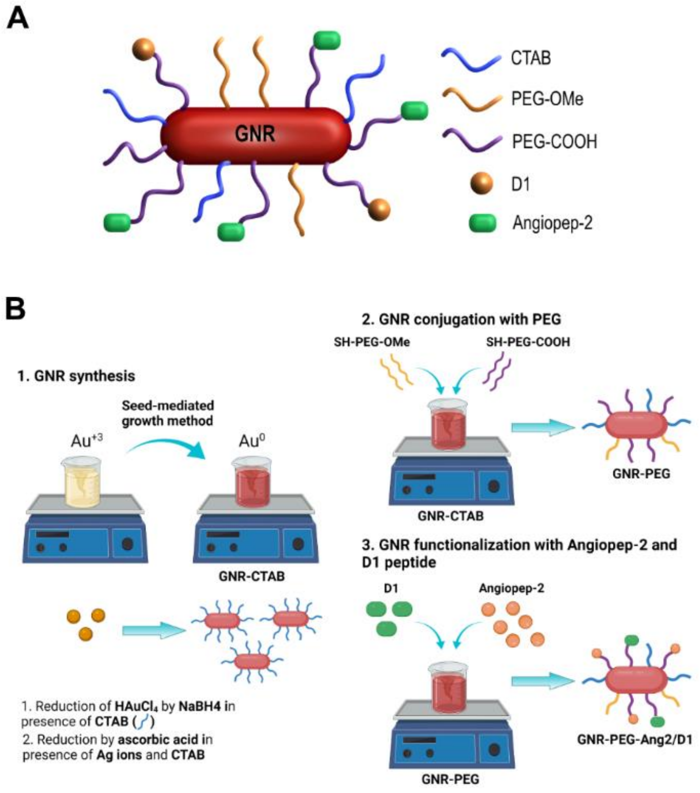


**Schema 2.** Schematic representation of A) GNR-PEG-Ang2/D1 and B) Synthesis and functionalization process.

A

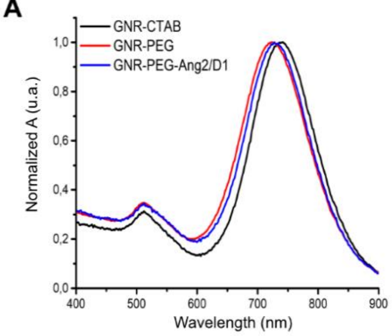


B

| Surface modification | Dh transversal (nm) ± SD | Dh longitudinal (nm) ± SD | Pdi | pZ (mV) ± SD |
|---|---|---|---|---|
| **GNR-CTAB** | 1.25 ± 0.1 | 55.00 ± 1.0 | 0.5 | 37.00 ± 2.0 |
| **GNR-PEG** | 4.00 ± 0.1 | 74.00 ± 2.0 | 0.5 | -32.00 ± 1.0 |
| **GNR-PEG-Ang2/D1** | 4.00 ± 0.1 | 61.00 ± 2.0 | 0.4 | -8.00 ± 2.0 |

C

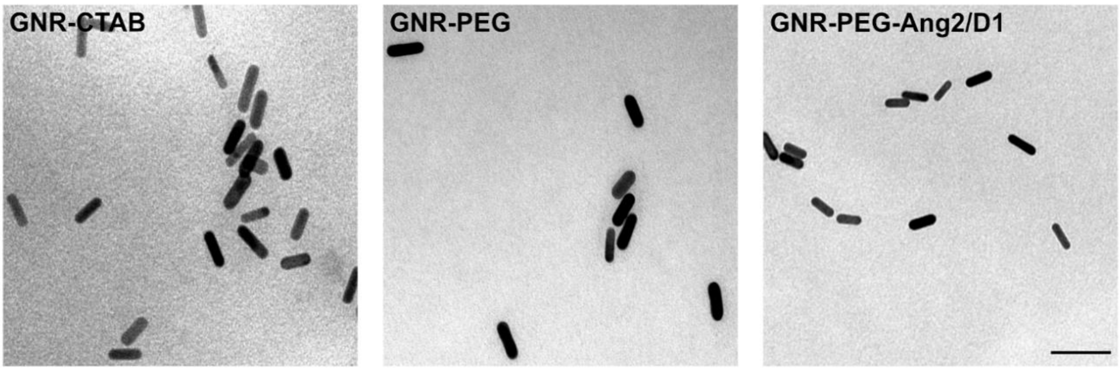


D

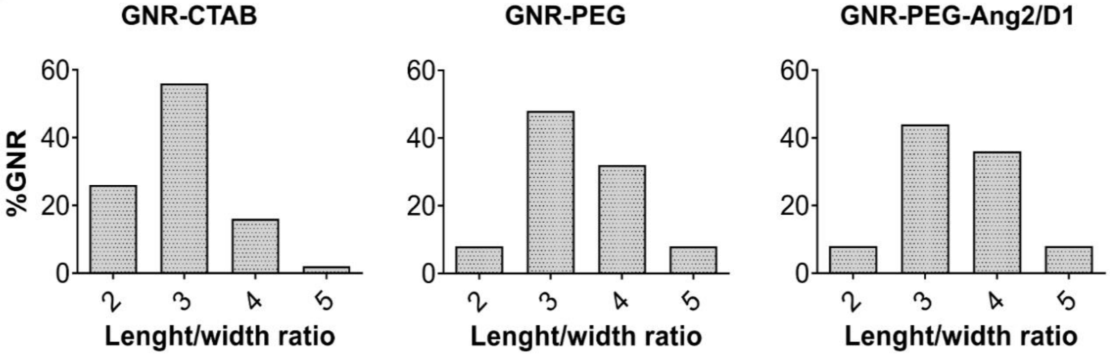


**Supplementary Figure S3.** Nanosystem characterization of GNR-CTAB, GNR-PEG and GNR-PEG-Ang2/D1. A) UV-vis-NIR absorbance spectrum B) Transversal and longitudinal hydrodynamic diameter (Dh), polydispersity index (Pdi) and zeta potential (pZ) C) TEM images obtained by transmission electron microscopy and D) histograms with their corresponding aspect ratio length/width distribution from 50 particles.

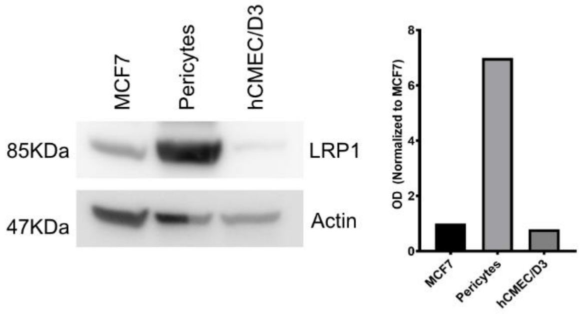


**Supplementary Figure S4.** Western blot of MCF7, pericytes and hCMEC/D3 cells lysate against LRP1 protein and actin.

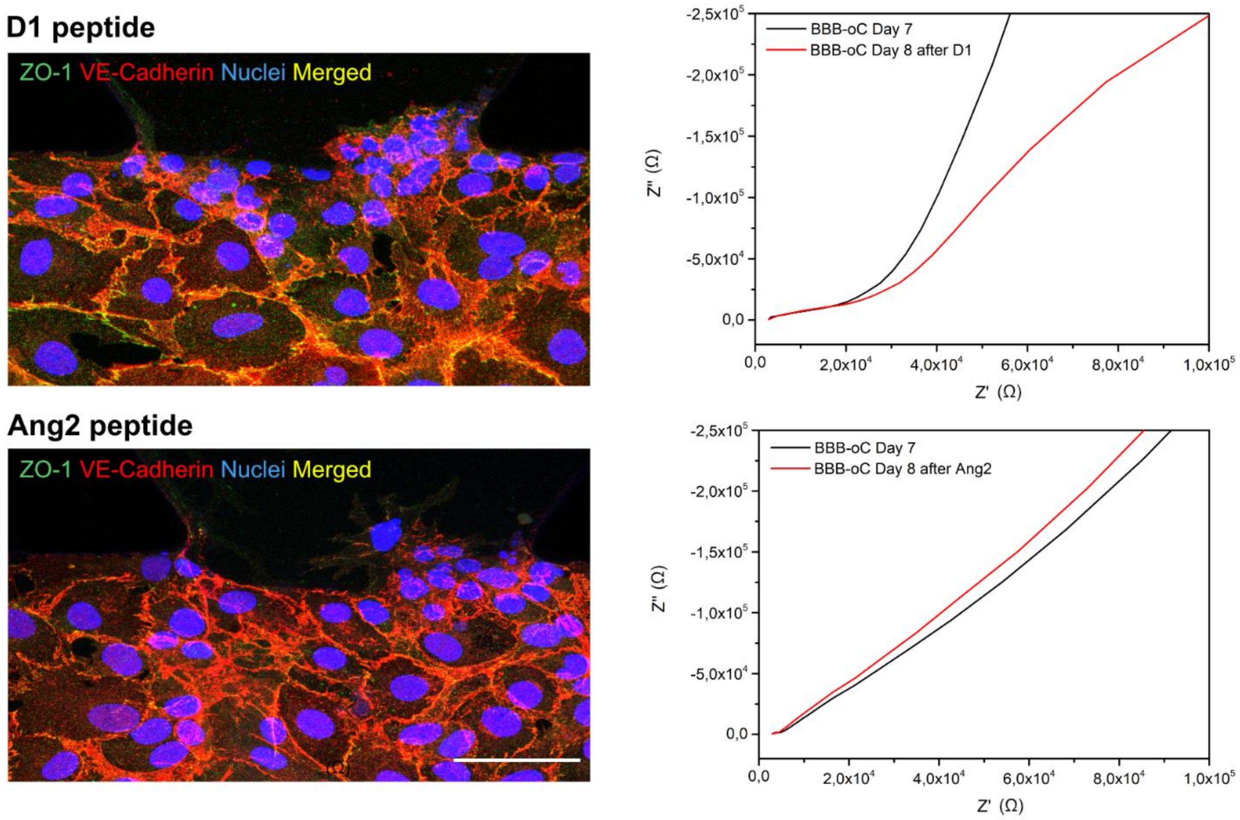


**Supplementary Figure S5.** TJs influence D1 and Ang2 peptides by immunofluorescence and TEER. Fluorescence confocal images of TJs on Day 8 after incubation with D1 175,6nM and Ang2 69,2nM for 24 hours. Scale bar 50µm. Nyquist plot of TEER measurement of BBB-oC before (Day 7) and next to peptide exposure for 24 hours.
